# Development and validation of a cellular host response test as an early diagnostic for sepsis

**DOI:** 10.1101/2021.02.01.429128

**Authors:** Lionel Guillou, Roya Sheybani, Anne E. Jensen, Dino Di Carlo, Terrell Caffery, Christopher Thomas, Ajay M. Shah, Henry T. K. Tse, Hollis R. O’Neal

## Abstract

Sepsis must be diagnosed quickly to avoid morbidity and mortality. However, the clinical manifestations of sepsis are highly variable and emergency department (ED) clinicians often must make rapid, impactful decisions before laboratory results are known. We previously developed a technique that allows the measurement of the biophysical properties of white blood cells as they are stretched through a microfluidic channel. In this study we describe and validate the resultant output as a model and score – the IntelliSep Index (ISI) – that aids in the diagnosis of sepsis in patients with suspected or confirmed infection from a single blood draw performed at the time of ED presentation. By applying this technique to a high acuity cohort with a 23.5% sepsis incidence (*n*=307), we defined specific metrics – the aspect ratio and visco-elastic inertial response – that are more sensitive than cell size or cell count in predicting disease severity. The final model was trained and cross-validated on the high acuity cohort, and the performance and generalizability of the model was evaluated on a separate low acuity cohort with a 6.4% sepsis incidence (*n*=94) and healthy donors (*n*=72). For easier clinical interpretation, the ISI is divided into three interpretation bands of Green, Yellow, and Red that correspond to increasing disease severity. The ISI agreed with the diagnosis established by retrospective physician adjudication, and accurately identified subjects with severe illness as measured by SOFA, APACHE-II, hospital-free days, and intensive care unit admission. Measured using routinely collected blood samples, with a short run-time and no requirement for patient or laboratory information, the ISI is well suited to aid ED clinicians in rapidly diagnosing sepsis.

## Introduction

Sepsis is a serious and potentially life-threatening condition, and one of the leading causes of death in the USA (1). Most patients with suspected infection who are at risk of sepsis will present to the hospital via the emergency department (ED), where clinicians must diagnose sepsis quickly and accurately in order to prioritize resources to care for those most at risk for clinical deterioration (2). Sepsis tends to progress very rapidly, and recent data suggests that each hour of delay to treatment results in an increase in morbidity and mortality (3, 4). The rapid and accurate diagnosis of sepsis is necessary for ED clinicians to quickly provide antibiotics and resuscitation to the most acutely ill patients, while limiting broad spectrum antibiotic use in less acutely-ill patients (5).

In practice, sepsis can be challenging to identify during an ED evaluation because the clinical manifestations of sepsis are highly variable (6). Clinical criteria for systemic inflammatory response syndrome, which previously served as a requisite for sepsis diagnosis, lack the sensitivity and specificity to accurately identify those patients most likely to progress to organ failure and death (7, 8). The Third International Consensus Task Force has defined sepsis as “life-threatening organ dysfunction due to a dysregulated host response to infection” and recommended that organ dysfunction be indicated by an increase in the Sequential Organ Failure Assessment (SOFA) score of 2 points or more (9). While this Sepsis-3 definition focuses on patients with higher mortality risk, clinical data and laboratory results required for the calculation of SOFA may be unavailable or difficult to obtain in a timely manner (8). As such, there is a critical need for a rapid test that can aid ED clinicians in identifying sepsis among patients with suspected or confirmed infection.

Inherent in the Sepsis-3 definition is the recognition that the patient’s immune system becomes dysregulated in response to an infection, leading to aberrant immune activation and subsequent clinical deterioration. In previous work, we developed a technique to rapidly quantify the state of immune activation by measuring the biophysical properties of white blood cells (WBCs) (10–12). By examining the physical changes that occur as cells are stretched through a microfluidic channel, this approach provides label-free, quantitative biophysical markers of immune activation. An earlier study demonstrated that this technique can rapidly characterize thousands of single cells, and measure biophysical changes in subpopulations of WBCs that are distinct in patients with sepsis (13).

In this study, we present the development and internal validation of a model and score – the IntelliSep Index (ISI) – that aids in the diagnosis of sepsis in patients with suspected or confirmed infection. The model was trained and cross-validated with data from a high acuity cohort, and data from a low acuity cohort and healthy volunteers were used to evaluate the generalizability of the model and estimate its expected clinical performance. The robust performance of the model indicates that the ISI may be useful as an aid to ED clinicians in the rapid and accurate diagnosis of sepsis.

## Material and methods

### Study design and setting

This work involved three prospective cohorts from separate studies – a high acuity cohort (*n*=307), a low acuity cohort (*n*=94), and a healthy donor cohort (*n*=72). The high acuity cohort was used for feature generation and selection, and as a training set to derive the final model reporting the ISI, whereas the low acuity cohort and the healthy donor cohort were used to evaluate feature and model performance on new datasets. Peripheral blood samples were collected from February 2016 to December 2016 for the high acuity cohort, and from July 2017 to January 2018 for the low acuity cohort, from subjects over 18 years of age who presented to the ED with signs suggestive of infection at two academic medical centers, namely Our Lady of the Lake Regional Medical Center and Baton Rouge General Medical Center, in Baton Rouge, LA. Subjects were identified for recruitment using the site’s electronic medical records system. Blood samples for the healthy donor cohort were provided by Our Lady of the Lake Blood Donor Center, Baton Rouge, LA, that collected blood from healthy volunteers who donated blood between June 2018 and January 2020, and completed a questionnaire indicating no current illnesses. All three studies were approved by the Louisiana State University Health Sciences Center Institutional Review Board (IRB) as well as by local, hospital-specific IRBs as appropriate (high acuity: LSUHSC-NO #8964, low acuity LSUHSC-NO #9749, healthy: FRANU #2018-022). The study team obtained written informed consent from all subjects. A detailed flow chart for how we selected evaluable subjects in each cohort is presented in S1 Fig.

The high acuity cohort enrolled subjects with two or more criteria for Systemic Inflammatory Response Syndrome (SIRS 2+) and at least one sign of potential organ dysfunction, defined as serum lactate > 2 mmol/L, altered mental status, hypoxia (peripheral capillary oxygen saturation < 90% on room air), hypotension (systolic blood pressure < 90 mmHg), acute kidney injury, total bilirubin > 2.5 mg/dL, platelet count < 100,000 cells/μL, international normalized ratio (INR) > 1.5 and no history of vitamin k antagonist use. The low acuity cohort enrolled subjects with SIRS 2+ with or without evidence of organ dysfunction. Of note, the low acuity cohort excluded abnormal WBC count from SIRS criteria in an effort to enroll subjects either before laboratory data were available or those who did not have laboratory studies performed.

Subjects with hematologic malignancies, myeloproliferative diseases, and neutropenia (absolute neutrophil count < 1000 cells /μL) were excluded. In addition, potential subjects who were undergoing chemotherapy or with late-stage cancer, and those receiving macrolide class antibiotics for greater than 3 hours prior to enrollment, were deemed non-evaluable and were excluded from analysis. These subjects were excluded because of the unknown potential effects of these conditions and medications on the biophysical properties of WBCs.

For all three studies, upon consent, an EDTA-anticoagulated peripheral blood sample was obtained from the subject for the IntelliSep test and testing was performed on samples within 3 hours of sample blood collection.

### Clinical classification of subjects

Pre-existing conditions and comorbidities were recorded using the subject’s medical record. Diagnosis of infection (such as pneumonia) was obtained through retrospective adjudication by physicians with access to the subject’s entire medical record, including results of tests, cultures, other lab tests per standard of care, and imaging results. For the determination of sepsis, two independent physician adjudicators reviewed the subject’s medical record to assess whether they had sepsis during the study as defined by the Sepsis-3 consensus guidelines (i.e., whether there is an infection present which caused a dysregulated immune response that led to organ dysfunction) (8). In cases of discordance, review of the subject’s medical record by a third physician adjudicator was used to make the final determination.

Measures for clinical assessment included sequential organ failure assessment (SOFA) scores (14), maximum over three days following ED presentation, acute physiology and chronic health evaluation (APACHE-II) scores (15), and predisposition insult response and organ failure (PIRO) scores (16). Utilization of hospital resources was measured by admission to the hospital, admission to the intensive care unit (ICU), in-hospital length of stay, excluding days deceased while in-hospital (LOS), and ICU LOS.

### IntelliSep test

Samples were evaluated using an early prototype version of the IntelliSep test (Cytovale, San Francisco, CA). A 100 µL sample of EDTA-anticoagulated whole blood was incubated with 5 µL PE-Cy5 CD45 (HI30 clone, Biolegend, cat. # 304010) and 5 µL of PE-CD66B (Becton Dickinson, cat. # 561650) for 5 minutes. Red blood cells present in the sample were lysed using the IMMUNOPREP Reagent System (Beckman Coulter) and red blood cell debris was washed away using the Lyse Wash Assistant (BD Biosciences), resulting in a total volume of 1 mL of WBCs suspended in PBS at a final concentration of 500-1000 cells/µL. The entire 1 mL of WBC suspension was transferred to a cartridge that was then inserted into an imaging module to interrogate single-cell biophysical properties via deformability cytometry (10–12). A controlled pressure of 90 pounds per square inch (psi) was applied to push the WBC suspension through the microfluidic channels, which contained an optical port to allow for high fidelity imaging at the junction where the cells were deformed. Data were collected capturing cells traveling through the microfluidic junction, yielding approximately 10-15 frames for each deforming cell. The number of cells analyzed for each subject was 47,000 +/−27,000 WBCs (mean +/-SD), and the run-time for each test using a prototype system was approximately 30 minutes end-to-end, including sample processing and image analysis.

Quality checks were performed for each test run to ensure that the instrument was performing as intended, and control objects (8.86 µm, PS-COOH beads, microParticles GmbH) were included in each sample to monitor the flow characteristics of the test run. Only test runs where the trajectory of the control objects was consistent with an unobstructed extensional flow region were accepted, while test runs in which the microfluidics were clogged were excluded. The velocity of the control objects in a straight channel upstream of the junction was also monitored to ensure that the target throughput was achieved. Other quality checks were performed in each test run to monitor any drift or vibration in the instrument, as well as poor lysis or wash performance during the sample preparation step.

### Image analysis and metrics

All the steps in the image analysis pipeline were automated, from the raw video all the way to the ISI, which was the final output. Images were analyzed by first passing the raw video (containing about 4 million frames) through an object detection algorithm to identify frames that contain cells. Images from those frames were then passed through a convolutional neural network that segmented each image to distinguish the cell from the background. Descriptors of cell shape, position, and morphology were computed for each image, and the metrics were aggregated at the event level, where an event indicates an instance of a single cell travelling through the extensional flow region. A primary metric, named visco-elastic inertial response (VEIR), measures cell oscillations, which are dependent on cell density, size, and visco-elastic properties. A secondary metric, termed aspect ratio (AR), quantifies the cell deformation under hydrodynamic stress fields, which reflects nuclear compressibility. For clarity, the term “metric” is used here to denote measures at the level of a single cell, whereas the term “feature” denotes measures at the level of cell populations.

WBC subpopulations (lymphocytes, neutrophils, and monocytes) were identified and enumerated by measuring cell size and optical intensity, using an automated clustering algorithm. The clustering algorithm was validated by comparing the cell clusters found by the automated clustering, to the fluorescent data obtained from PE-Cy5 CD45 and PE-CD66B stains using conventional flow cytometry techniques.

Once the metrics for each cell event were computed, and the cells segmented into subpopulations, statistical descriptors were computed to summarize the metrics into features. Some example features include the average size of neutrophils and the 75^th^ percentile of the VEIR of monocytes. Finally, the ISI was computed by linearly combining all the features. Next, we explain how the feature coefficients were determined during model training.

### Model training and validation

Only data from the high acuity cohort were used for model training. A cross-validation framework was used to train a linear logistic classifier to disease state (septic or not septic), using features as inputs. For classification, a logistic regression with a L1 regularization available through scikit-learn’s sklearn.linear_model package was used. Scikit-learn is a free and open-source Python library for data analysis and machine learning applications (17). To guard against over-fitting, data from approximately 25% of subjects in the high acuity cohort were held out as a sequestered set for validation testing (sequestered testing set, *n*=69). All of the feature selection and model training was performed on a k-fold nested cross-validation architecture on data from the remaining 75% of subjects (initial training set, *n*=238). Once the features were finalized, the model was validated using the sequester set to confirm the generalizability of the model development methodology.

The final model was generated by integrating the sequestered testing set into the initial training set for a final training step, which included data from all of the subjects in the high acuity cohort (full training set, *n*=307). The ISI was produced by linearly transforming the prediction from the final model in the log-odds space into a single score between 0.1 and 10.0, inclusive. To allow for easier clinical interpretation, the range of the scores are divided into three interpretation bands of Green (0.1 – 5.4), Yellow (5.5 – 6.7), and Red (6.8 – 10.0) that correspond to increasing probability of sepsis. The cutoffs for the interpretation bands were established based on observations from the full training set. Predictions of ISI using data from the low acuity cohort and the healthy donor cohort were used to assess the performance and generalizability of the final model.

### Statistical analysis

Unless otherwise stated, p-values were obtained from an unpaired two-sample Welch’s t-test or Mann-Whitney U test, as appropriate, where the null hypothesis is that the mean of the two samples are equal. Significance levels are indicated throughout the manuscript as follows: * p < 0.05, ** p < 0.01, *** p < 0.001. Descriptive statistics are presented as means, standard deviations, medians, and interquartile ranges for the continuous variables, and as counts and percentages for categorical variables. An alpha level of 0.05 was used for all analyses, unless otherwise stated. Two-sided confidence intervals for proportions are provided using the Clopper-earson method, where appropriate. Two-sided confidence intervals for ROC AUC were computed using a bootstrap method performed with 1000 replicates, where model predictions on the test set were resampled with replacement to estimate a probability distribution for the AUC. Time-to-event analyses are depicted with Kaplan-Meier plots, and the log-rank test was used for comparison between groups.

## Results

### Subject demographics

The three separate prospective cohorts used for this work – a high acuity cohort, a low acuity cohort, and a healthy donor cohort – represented a broad spectrum of potential disease severity. Baseline characteristics of the evaluable subjects from all three cohorts are presented in Table 1. For the high acuity cohort, the median age was 62 years (Q1-Q3 50-74), with 153 male (49.8%) and 154 female (50.2%). Of the total, 53.1% were White, 40.1% were African American, and 6.8% were members of other races. Median age was significantly lower for the low acuity and healthy study populations at 55 (Q1-Q3 42 – 71) and 56 (Q1-Q3 38 – 68), respectively. Sex and race distributions for the low acuity and healthy studies were similar to the high acuity study, except for the significantly higher representation of While individuals in the healthy cohort. As expected, the high acuity cohort had the highest rate of sepsis (23.5%) as adjudicated by consensus retrospective physician review, which was significantly higher than the 6% sepsis rate among the low acuity cohort (p< 0.001). The high acuity cohort also utilized more hospital resources (hospital admission, hospital length of stay, ICU admission), and had greater severity of illness (higher SOFA, APACHE-II, and PIRO scores) and higher mortality as compared to the low acuity cohort.

**Table 1.** Baseline characteristics of subjects in each of the three cohorts.

| Characteristic | High Acuity Cohort<br>N = 307 | Low Acuity Cohort<br>N = 94 | Healthy Donor Cohort<br>N = 72 | p-value(s) |
| --- | --- | --- | --- | --- |
| Age, median (IQR) | 62.2 (50.4, 73.6) *, † | 55.3 (42.3, 70.6) * | 52.5 (38.0, 68.0) † | * $p < 0.01$<br>† $p < 0.001$ |
| Sex, Female, N (%) | 154 (50.2) | 44 (46.8) | 40 (55.6) | NS |
| Race, N (%) |  |  |  |  |
| White | 163 (53.1) * | 47 (50.0) † | 63 (87.5) *, † | * $p < 0.001$<br>† $p < 0.001$ |
| African American | 123 (40.1) * | 44 (44.8) † | 4 (5.6) *, † |  |
| Other | 21 (6.8) | 3 (3.2) | 3 (4.2) |  |
| Infection Source, N (%) |  |  |  |  |
| Pulmonary | 37 (12.0) | 7 (7.4) | N/A | NS |
| Genitourinary | 24 (7.8) | 4 (4.3) | N/A |  |
| Skin-Skin Structure / Ortho / Wound | 24 (7.8) | 1 (1.1) | N/A |  |
| Abdominal / Gastrointestinal | 23 (7.5) | 4 (4.3) | N/A |  |
| Primary Bloodstream | 9 (2.9) | 2 (2.1) | N/A |  |
| Central Nervous System | 1 (0.3) | 0 (0.0) | N/A |  |
| Not Infected | 197 (64.2) *, † | 76 (80.9) *, ‡ | 72 (100) ‡, † | * p < 0.001<br>† p < 0.001<br>‡ p < 0.001 |
| Classification: Consensus RPD, N (%) |  |  |  |  |
| Sepsis | 72 (23.5) * | 6 (6.4) * | N/A | * p < 0.001 |
| SIRS/Not Sepsis | 235 (76.5) * | 88 (93.6) * | N/A |  |
| Admitted to Hospital, N (%) | 293 (95.4) * | 65 (69.1) * | N/A | * p < 0.001 |
| Admitted to ICU, N (%) | 81 (26.4) * | 11 (11.7) * | N/A | * p < 0.001 |
| Hospital LOS (days), median (IQR) | 4.0 (2.0, 6.0) * | 2.0 (0.0, 4.0) * | N/A | * p < 0.001 |
| ICU LOS (days), median (IQR) | 0.0 (0.0, 1.0) | 0.0 (0.0, 0.0) | N/A | NS |
| In-Hospital Mortality, N (%) | 31 (10.1) * | 3 (3.2) * | N/A | * p < 0.01 |
| SOFA, 3-day max, median (IQR) | 4 (2, 5) * | 2 (0, 3) * | N/A | * p < 0.001 |
| APACHE-II, median (IQR) | 15 (12, 19) * | 11 (8, 14) * | N/A | * p < 0.001 |
| PIRO, median (IQR) | 5 (2, 7) * | 3 (1, 4) * | N/A | * p < 0.001 |
| WBC (10<sup>3</sup> cells/μL), median (IQR) | 13.8 (10.1, 17.5) *, † | 10.3 (7.5, 13.0) *, ‡ | 6.0 (4.9, 7.0) ‡, † | * p < 0.001<br>† p < 0.001<br>‡ p < 0.001 |
| Triage Temperature, median (IQR) | 98.2 (97.8, 98.7) | 98.3 (98.0, 98.8) | N/A | NS |
| Triage Heart Rate, median (IQR) | 94.0 (81, 106) | 99.5 (88, 109) | N/A | NS |
| Triage Respiratory Rate, median (IQR) | 19 (17, 22) | 20 (16, 23) | N/A | NS |
| Lactate Measured, N (%) | 212 (69.1) * | 36 (38.3) * | N/A | * p < 0.001 |
| Lactate, median (IQR) | 2.4 (1.5, 3.6) | 1.9 (1.2, 2.9) | N/A | NS |
| IntelliSep Index, median (IQR) | 4.8 (3.7, 6.1) *, † | 4.3 (3.4, 5.2) *, ‡ | 2.2 (1.5, 3.2) ‡, † | * p < 0.01<br>† p < 0.001<br>‡ p < 0.001 |

Abbreviations: APACHE-II, acute physiology and chronic health evaluation; ICU, intensive care unit; IQR, interquartile range; LOS, length of stay; N/A, not applicable; NS, not significant; PIRO, predisposition infection response organ; RPD, retrospective physician diagnosis; SOFA, sequential organ failure assessment; SIRS, systemic inflammatory response syndrome; WBC, white blood cell count. p-values were obtained from an unpaired two-sample Welch's t-test, with the null hypothesis that the mean of the two samples are equal.

### Feature performance

Representative scatter plots of cell size as compared to AR and VEIR for a healthy donor, a subject adjudicated as non-septic SIRS 2+, and a subject adjudicated as septic are shown in Fig 1. For ease of interpretation, cells with high AR and high VEIR across all the scatters are highlighted in red. The data show an increasing number of cells with high AR and high VEIR corresponding to the adjudicated disease state of the subject – from healthy, to non-septic SIRS 2+, to septic – indicating that AR and VEIR may characterize the immune activation state of the measured leukocytes. Interestingly, cell size is not correlated with either AR or VEIR, suggesting that additional information is provided by these measures. The data also show that higher AR and higher VEIR lead to higher ISI values, and suggest that higher ISI values correlate with increasing levels of immune activation.

**Fig 1.**
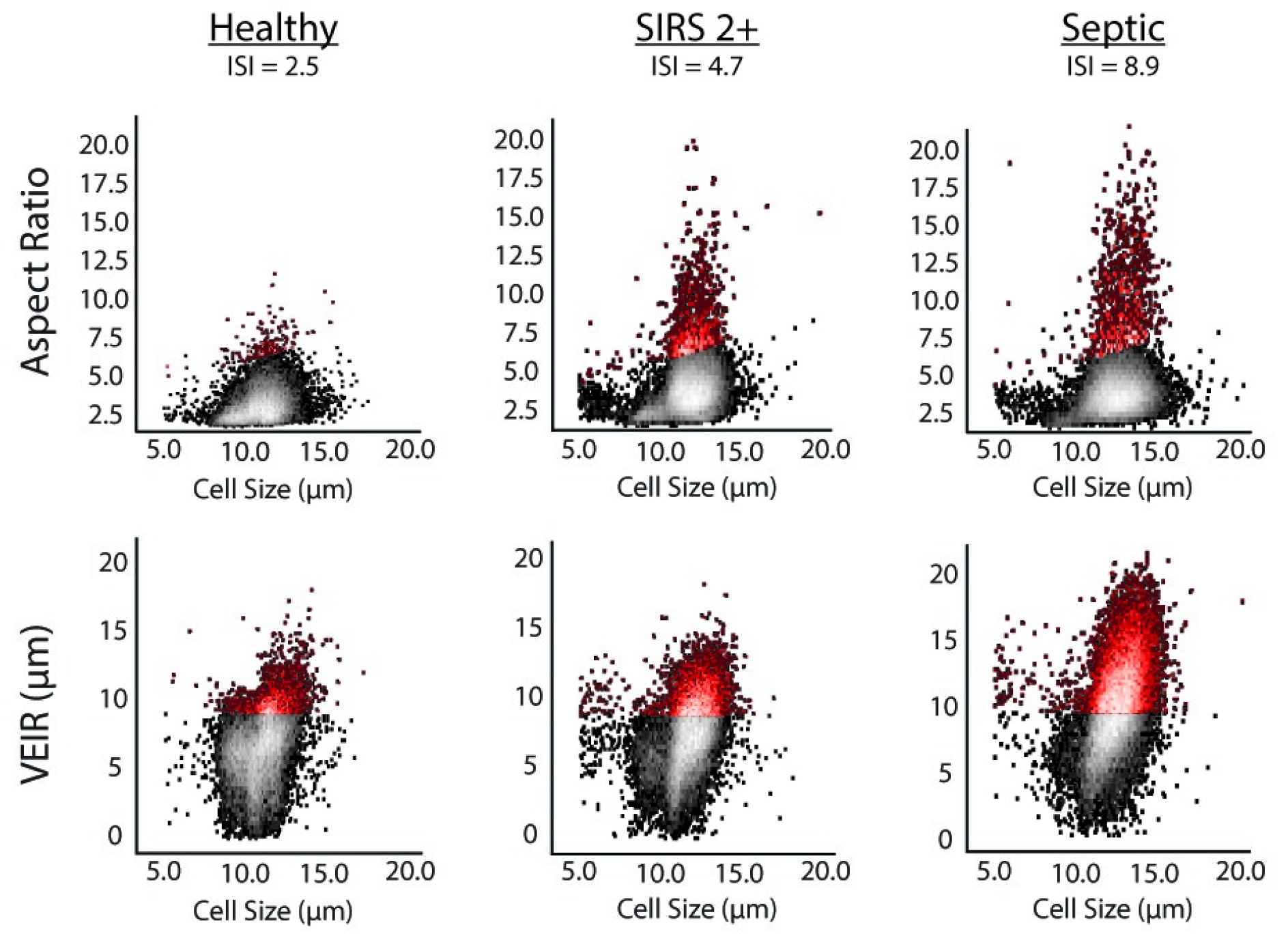
**Representative scatter plots of cell size versus aspect ratio and visco-elastic inertial focusing (VEIR) from a healthy donor, a subject adjudicated as non-septic SIRS 2+ from the low acuity cohort, and a subject adjudicated as septic from the high acuity cohort.** As a visual aid, cells with high AR and high VEIR are hued in red. For the purposes of this plot only, the cutoff for “high AR” was arbitrarily chosen as all the points defined by AR ≥ 0.3 + 2.5 x (Cell Size), and the cutoff for “high VEIR” was arbitrarily chosen as 9 µm.

The performance of features in subjects with increasing severity of disease were further explored by evaluating cell count, cell size, AR and VEIR for the entire population of healthy donors as compared to subjects with SIRS 2+ and subjects with sepsis (Fig 2). The data show statistically significant increases in cell count, cell size, AR, and VEIR of neutrophils and monocytes for subjects with symptoms associated with increased immune activation. Specifically, mean neutrophil AR increased from 2.55 ± 0.24 (median ± SD) for the healthy donor cohort to 3.13 ± 0.32 for septic subjects, and mean monocyte AR from 3.24 ± 0.37 to 3.42 ± 0.60, while mean neutrophil VEIR increased from 6.73 ± 0.60 µm to 8.19 ± 0.87 µm, and mean monocyte VEIR from 8.26 ± 0.79 µm to 10.41 ± 0.90 µm. Trends for lymphocytes, while statistically significant, appeared to be much weaker than those for neutrophils and monocytes. Of note, cell size was not as sensitive as AR and VEIR in distinguishing disease status. This was particularly true for monocytes, where cell size was not statistically different between healthy donors, subjects with SIRS 2+, or septic subjects, and hence was not as useful as AR and VEIR in distinguishing sepsis. These observations are further supported by the effect size as measured by Cohen’s d, shown in S1 Table.

**Fig 2.**
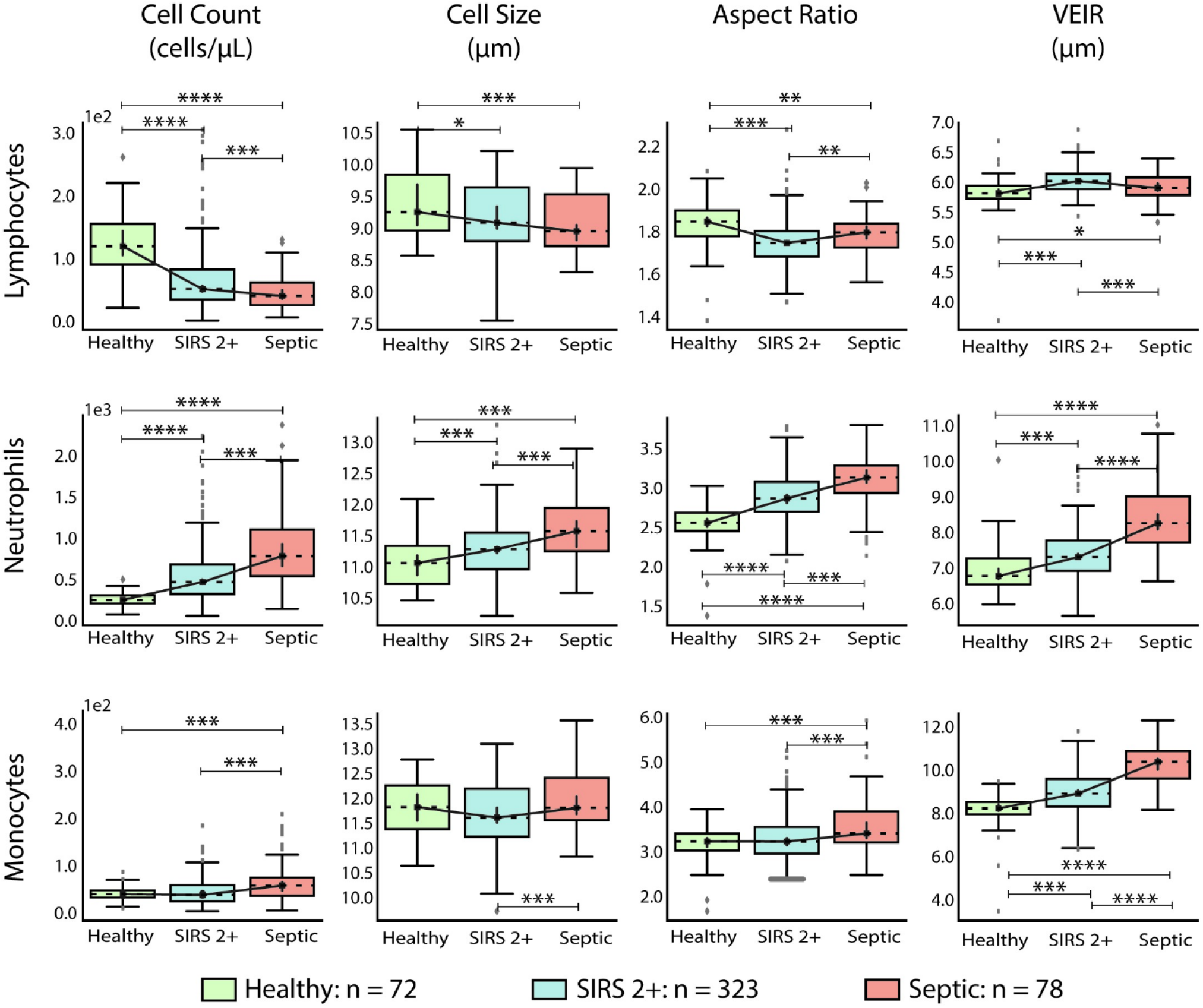
**Box plots displaying cell count, cell size, AR, and VEIR from healthy donors (*n*=72), subjects with non-septic SIRS 2+ (*n*=323), and subjects with sepsis (*n*=78), with features stratified by cell type (lymphocytes, neutrophils, and monocytes).** For these experiments, data from subjects presenting to the ED with 2 or more criteria for SIRS who were later adjudicated as non-septic or as septic were drawn from both the high acuity and low acuity cohorts. Mean values were computed for each cell type within a single test run. A p-value < 0.001 was observed using one-way analysis of variance for each feature for each cell type across the three cohorts, except for mean cell size for lymphocytes, where p-value was 0.002. The p-value from a post-hoc unpaired Student’s two-sided t-test comparing two columns at a time are indicated as follows: * p < 0.05, ** p < 0.01, *** p < 0.001, **** p < 10^−10^. Box-plot elements: center line, median; box limits, upper and lower quartiles; whiskers, 1.5x interquartile range; points, outliers.

Feature performance was further explored in subsets of healthy donors, subjects with SIRS 2+ and septic subjects that had normal WBC counts of 4×10^6^ to 12×10^6^ cells/mL (Fig 3). Within this subset of subjects, WBC count alone, as is commonly provided by a complete blood count (CBC), provides no indication of sepsis status. By comparison, both AR and VEIR are markedly higher in septic subjects with normal WBC counts. VEIR in particular is able to distinguish septic subjects, yielding values that are higher by approximately 2 µm for both neutrophils and monocytes, corresponding to a Cohen’s d larger than 2 for both cell types, showing a large effect size. These results further support that additional information is provided by AR and VEIR as compared to traditional cell counts.

**Fig 3.**
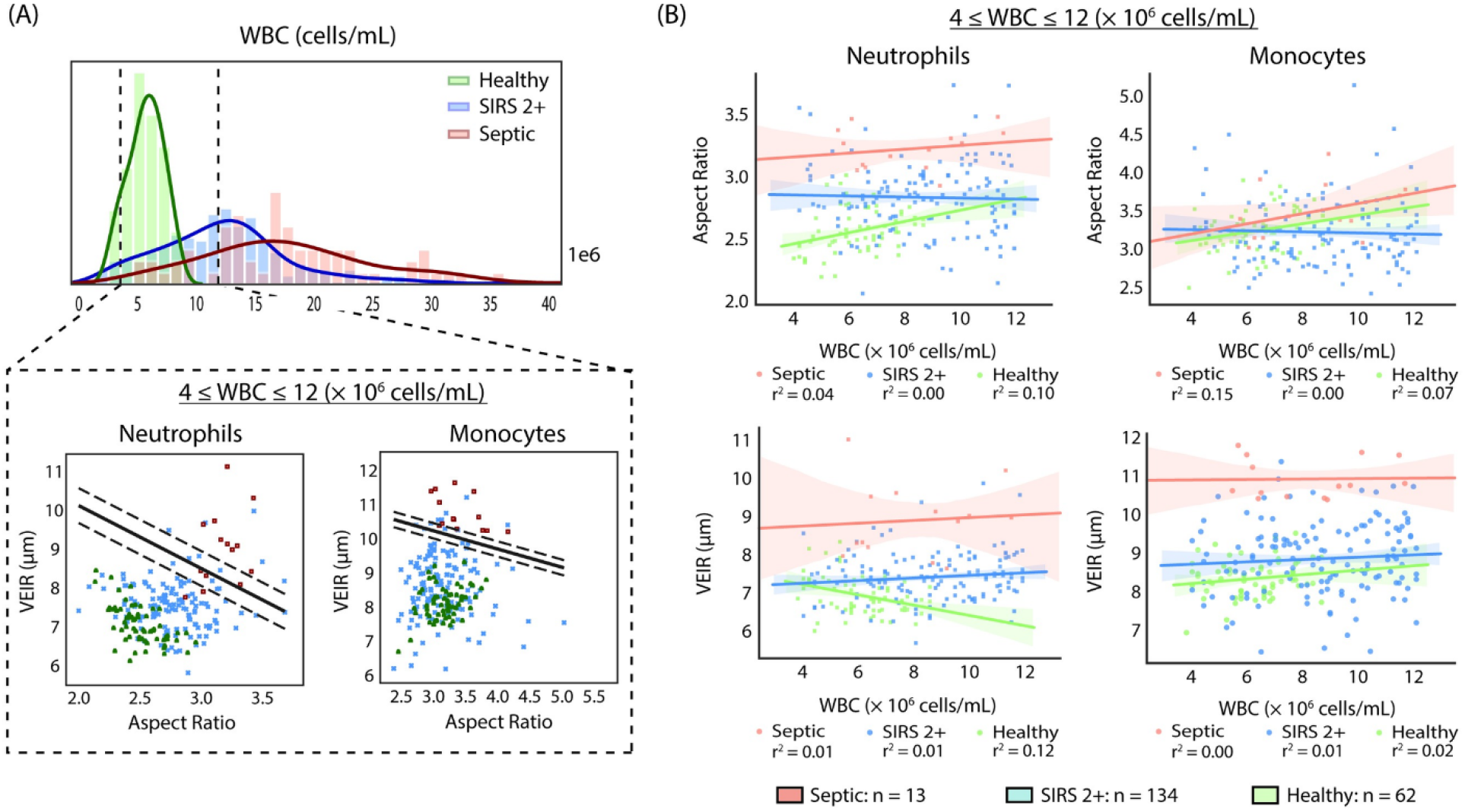
**(A, top panel) Histogram of WBC counts in healthy donors (*n*=72), subjects with SIRS 2+ (*n*=323), and subjects with sepsis (*n*=78). (A, bottom panel) Scatter plots of VEIR and Aspect Ratio in neutrophils and monocytes from the subset of healthy donors (*n*=62), subjects with SIRS 2+ (*n*=134) and subjects with sepsis (*n*=13) that have normal WBC counts between 4 × 10^6^ and 12 × 10^6^ cells/mL**. The lines on each graph indicate the decision boundary and the margins for a linear Support Vector Machine classification algorithm that is able to separate subjects between septic and non-septic using mean VEIR and mean AR features. (B) Scatter plots of AR and VEIR in neutrophils and monocytes from subjects with normal WBC counts. The lines indicate the linear regression between Aspect Ratio or VEIR and cell count.

### Model performance

The performance of the ISI, which is produced by linearly transforming the prediction in the log-odds space from the final model, was evaluated in a series of cross validation experiments. As described more fully in Materials and Methods, only data from the high acuity cohort was used to develop the final model, using an initial training set (*n*=238), a sequestered testing set (*n*=69), and then the full training set (*n*=307).

A first evaluation of model performance was conducted using a cross-validation framework with the initial training set, resulting in an area under the receiving operating curve (AUC) of 0.92 (0.87 – 0.96; 95% CI). To further test generalizability, the model trained on the initial training set was tested on the sequestered testing set, resulting in an AUC of 0.90 (0.80 – 0.98; 95% CI). The final model was then trained on the full training set with an AUC of 0.91 (0.87 – 0.95; 95% CI) (Fig 4A).

**Fig 4.**
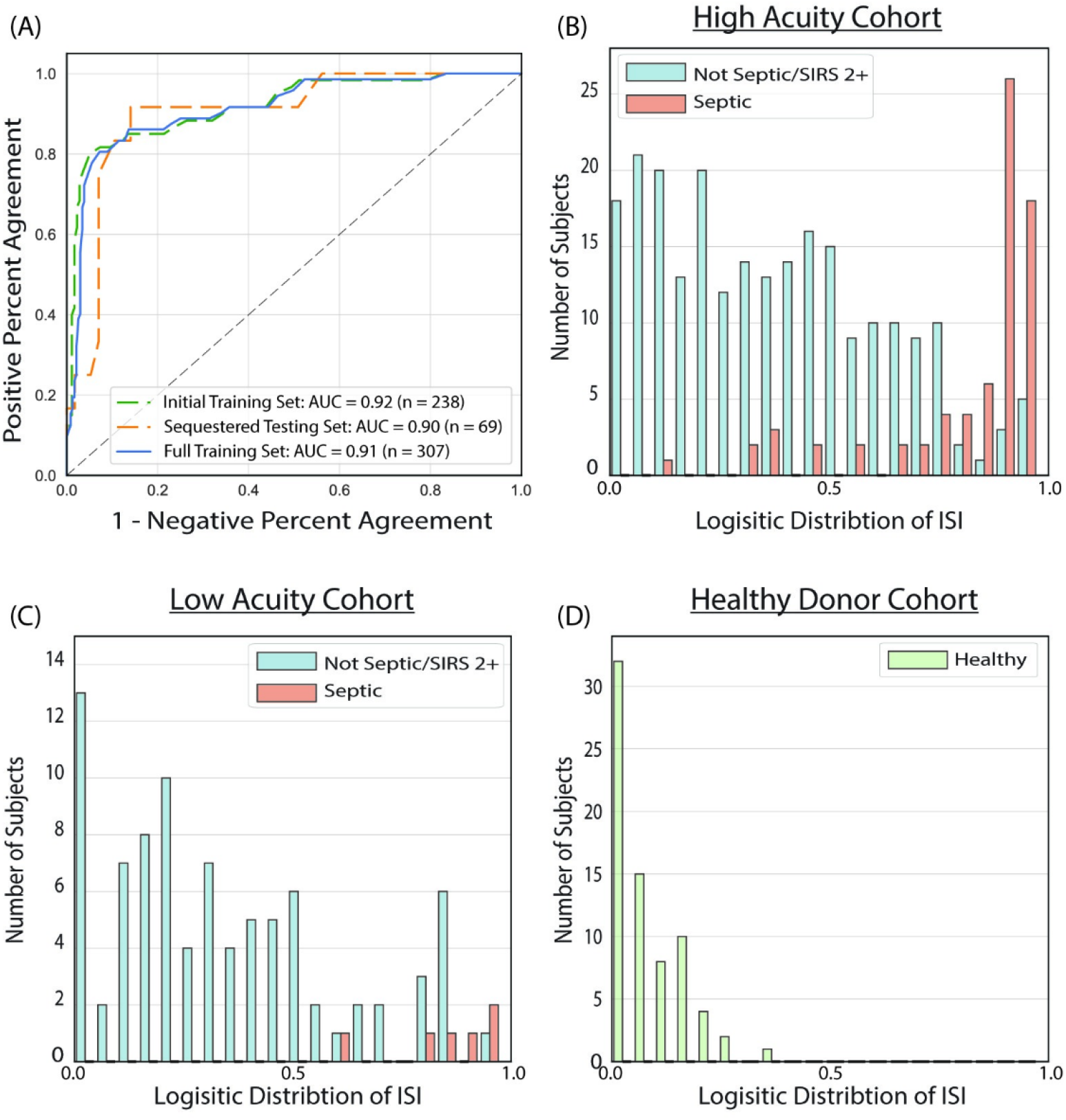
**(A) Receiver Operating Characteristic (ROC) curves showing the performance of the ISI as a diagnostic binary classifier to distinguish septic vs. non-septic subjects for the initial training set (*n*=238, adjudicated sepsis incidence of 25.2%), the sequestered testing set (*n*=69, adjudicated sepsis incidence of 17.4%), and the full training set (*n*=307, sepsis incidence of 23.5%). Also shown are logistic distributions of ISI values for (B) the high acuity cohort (*n*=307, adjudicated sepsis incidence of 23.5%), (C) the low acuity cohort (*n*=94, sepsis incidence of 6.4%), and (D) the cohort of healthy donors (*n*=72).**

In a separate set of cross validation experiments, the high acuity cohort was randomly partitioned into three folds, where models were trained on data from 2/3 of the data (*n*=205) and tested on the remaining 1/3 (*n*=102), using the same method used to create the final model. AUC values derived for these models were 0.89 (0.82 – 0.95; 95% CI), 0.90 (0.79 – 0.98; 95% CI) and 0.93 (0.86 – 0.98; 95% CI), which were close to the AUC of the final model of 0.91 (0.81 – 0.96; 95% CI), as shown in S2 Fig. The high acuity cohort was also randomly partitioned 100 times into 10 folds, where models were trained on 9/10 of the data and tested on the remaining 1/10 of the data. As shown in S3 Fig, the mean AUC from the repeated 10-fold partitions was 0.91, which again was close to the AUC found for the sequester set as well as the final model. Taken together, these results indicate that the performance of the model was not the result of overfitting, and would generalize well beyond the high acuity cohort.

Data from separate cohorts (low acuity and healthy population) with different incidence rates of sepsis were evaluated to assess the generalizability of the model and estimate its expected clinical performance. The data show that the ISI distinguished septic from non-septic subjects in the high acuity cohort (Fig 4B), as well as the low acuity cohort (Fig 4C) and the cohort of healthy donors (Fig 4D). In all three cohorts, septic subjects had higher ISI values that were clearly distinct from lower ISI values of the non-septic subjects.

In the high acuity cohort, 195 (63.5%) of subjects were in the Green Band, 53 (17.3%) in the Yellow Band, and 59 (19.2%) in the Red Band. A negative predictive value for sepsis (NPV) of 95% (83 – 98, 95% CI) for subjects in the Green Band and a diagnostic odds ratio of 102 when comparing those in Green and Red Bands was observed. Similar performance trends were observed for the low acuity cohort. As expected, a higher proportion of subjects fell in the Green Band (73, 77.7%), compared to 10 (10.6%) in the Yellow Band, and 11 (11.7%) in the Red Band. The observed NPV was 100% (40 – 100, 95%CI) for subjects in the Green Band. Note that the wide 95% CIs reported for the low acuity cohort is due to the low number of septic patients. All healthy donors had a low ISI value that fell within the Green Band. There were no significant differences in baseline demographics (age, sex, and race) across interpretation bands for any of the cohorts (S2 Table).

### Clinical studies

Table 2 shows the diagnosis for subjects in the high acuity cohort as determined by retrospective physician adjudication, separated into those with high ISI (Red Band) and those with low ISI (Green Band). The data show that not all subjects in the Red Band were adjudicated as having sepsis (right side of Table 2). However, nearly 90% of subjects in the Red Band had a diagnosis associated with activation of the immune system (i.e., either sepsis or a non-infectious severe immune response). By contrast, only 5% of subjects in the Green Band were adjudicated as septic (left side of Table 2), and the rest were diagnosed as having conditions not expected to be associated with activation of the immune system.

**Table 2.** Diagnosis of subjects in the high acuity cohort separated into those with high ISI (Red Band) and those with low ISI (Green Band).

| Green Band (ISI 0.1 – 5.4)<br>N = 195 |  | Red Band (ISI 6.8 – 10.0)<br>N = 59 |  |
| --- | --- | --- | --- |
| Congestive Heart Failure/ Chest Pain/Atrial Fibrillation | 54<br>(27.7%) | Sepsis | 50<br>(84.7%) |
| Chronic Respiratory Disease/COPD | 39 (20.0%) | Non-infectious Severe Immune Response (flare of Ulcerative Colitis, Rheumatoid Arthritis Pericarditis, Sarcoidosis) | 3<br>(5.1%) |
| Abdominal Pain/GI Bleed/Volume Depletion | 19<br>(9.7%) | Infection with Chronic/Direct Organ Dysfunction | 2<br>(3.4%) |
| Complications of Diabetes | 16<br>(8.2%) | Cardiac Arrest/Emergency Heart Failure | 2<br>(3.4%) |
| Infection without Organ Dysfunction | 14<br>(7.2%) | Other | 2<br>(3.4%) |
| Alcohol/Substance Abuse | 11<br>(5.6%) |  |  |
| Chronic Kidney Disease/ESRD | 10<br>(5.1%) |  |  |
| Encephalopathy/seizures | 10<br>(5.1%) |  |  |
| Sepsis | 10<br>(5.1%) |  |  |
| Chronic liver Disease | 4<br>(2.1%) |  |  |
| Direct Limb Injury/Fall | 4<br>(2.1%) |  |  |
| Other | 4<br>(2.1%) |  |  |

Trends in the severity of illness across the ISI interpretation bands are shown in Fig 5. For this analysis, the high acuity and low acuity cohorts were combined and subjects were evaluated by four measures, including SOFA score (maximum over 3 days), APACHE-II score, number of hospital-free days, and admission to the ICU. Hospital-free days were calculated as 28 days less the in-hospital LOS, where in-hospital mortality leads to a value of zero. The data show statistically significant differences between subjects in the Red Band, Yellow Band, and Green Band for all four measures, indicating that the ISI interpretation bands stratify for severity of illness and use of hospital resources.

**Fig 5.**
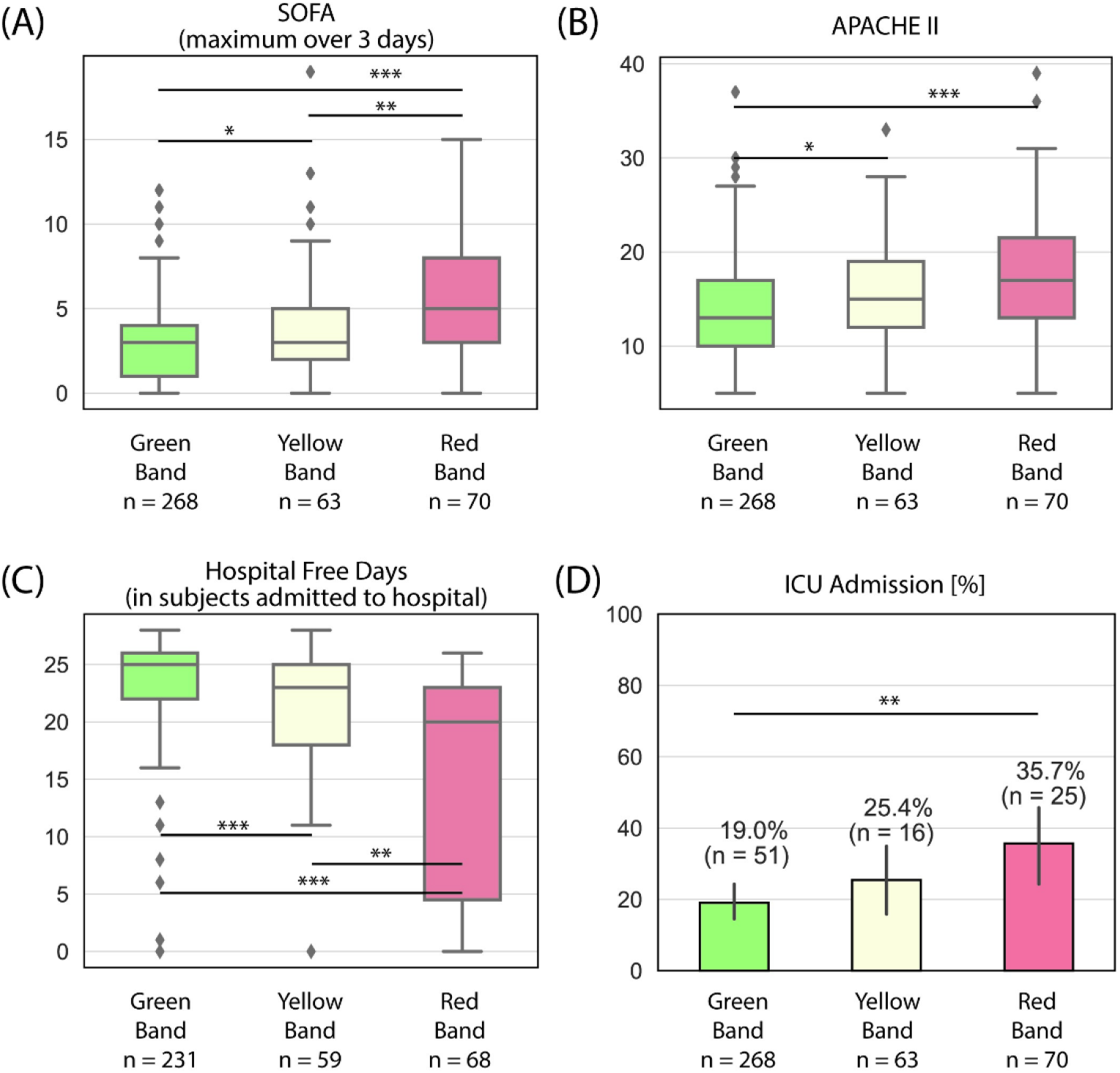
**Trends in severity of illness across ISI interpretation bands for subjects in the high and low acuity cohorts, as measured by (A) SOFA, maximum over three days following ED presentation, (B) APACHE-II, (C) hospital free days for the subpopulation of subjects admitted to the hospital, (D) Admission to the ICU.** Hospital free days were defined as 28 days less the in-hospital length of stay, where in-hospital mortality leads to a value of zero. Box plots: lines in the boxes, medians; the box ends, interquartile ranges (IQR); whiskers, 1.5x IQR; diamonds, outliers. Bar graphs: Bars, percentages; error pars, 95% confidence intervals. p-values were obtained from an unpaired two-sample Welch’s t-test (except for hospital free days, where the Mann–Whitney U was due to the non-normal distribution), with the null hypothesis that the mean of the two samples are equal. p-values reported as * p < 0.05, ** p < 0.01, and *** p < 0.001.

The potential clinical relevance of the ISI was further explored by Kaplan-Meier survival analysis of all subjects in the high acuity cohort (*n*=307) as well as the subpopulation of subjects adjudicated as infected by independent physician review (*n*=102). The data in Fig 6 show that the ISI interpretation band is correlated with survival probability. Specifically, subjects in the Red Band were clearly distinguished from those in the Green Band (p < 0.001, Fig 6A). Consistent with this result, among subjects adjudicated as infected, those in the Red Band were clearly distinguished from those in the Green Band (p < 0.01, Fig 6B). The survival curves for the subpopulation of infected subjects is notable in that all of these subjects presented to the ED with two or more criteria for SIRS and at least one sign of potential organ dysfunction. As such, the Kaplan-Meier analysis shown in Fig 6B reflects mortality from sepsis.

**Fig 6.**
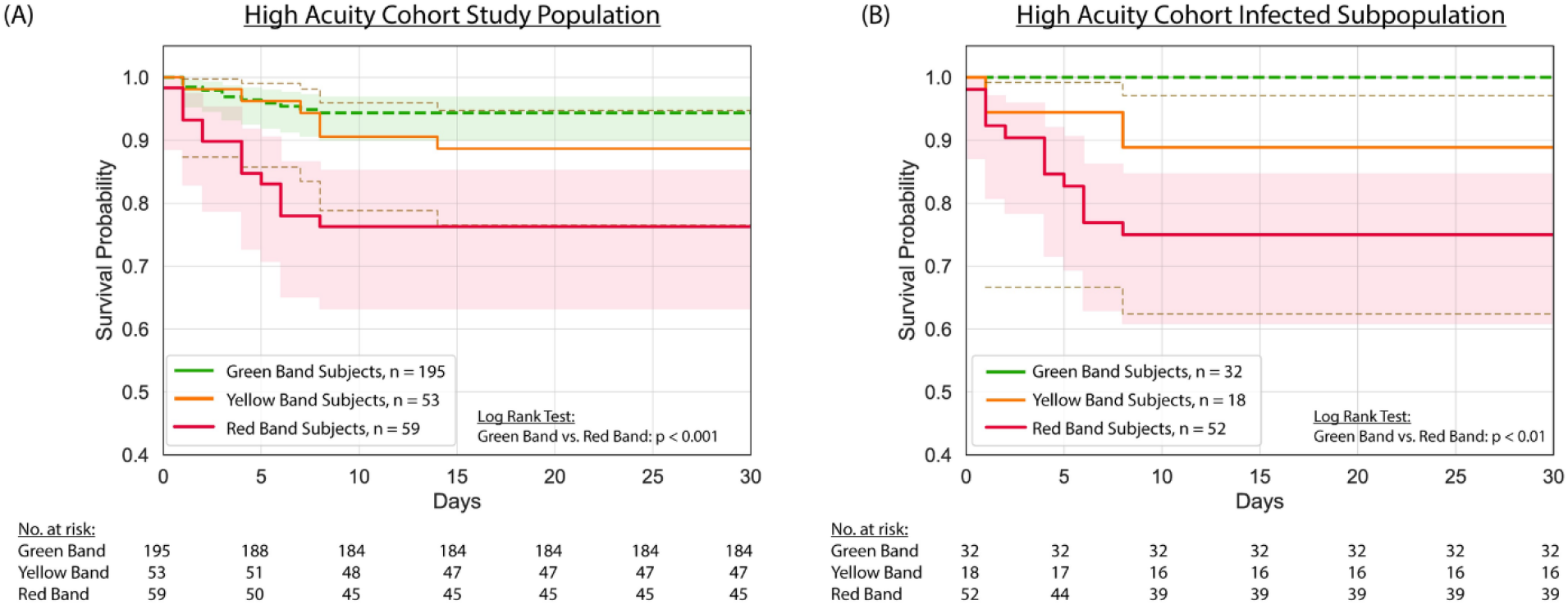
Kaplan-Meier survival stratified by ISI interpretation band for (A) all evaluable subjects (*n*=307) and (B) a subpopulation of subjects adjudicated as infected (*n*=102) in the high acuity cohort. Plots were created with right-censoring, with the assumption that subjects discharged from the ED or hospital survived ≥ 30 days in the absence of evidence to the contrary (e.g., return to the ED, discharge to hospice, or other indication in the electronic health record, which was reviewed after 30 days, that the subject died). Shading depicts 95% confidence intervals.

## Discussion

Early recognition and prompt treatment of sepsis is critical for reducing morbidity and mortality. Although time is of the essence, ED clinicians lack rapid, efficient tests to identify sepsis among patients with signs of infection. In this study, we present the development and initial validation of the ISI that may be useful as an aid for ED clinicians in the rapid and accurate diagnosis of sepsis. The ISI’s high NPV may enable ED clinicians to quickly identify and treat septic patients, while distinguishing non-septic patients for whom aggressive antibiotic therapy and resuscitation may not be warranted, and where an alternate diagnosis should be considered.

We developed the ISI using a technique that measures the biophysical properties of WBCs as they are stretched through a microfluidic channel (10–12). This technique enables the characterization of thousands of single cells in a matter of minutes, and provides information on the host immune system that can be used to distinguish patients with sepsis (13). By applying this technique to a high acuity patient population in the ED context, we were able to define specific metrics – the AR and VEIR – that are much more sensitive than measures such as cell size or cell count in predicting severity of disease. Our final model incorporates AR and VEIR to yield predictions on disease severity from which the ISI is derived. For ease of clinical interpretation, we divided the ISI into three interpretation bands of Green (0.1 – 5.4), Yellow (5.5 – 6.7), and Red (6.8 – 10.0) that correspond to increasing disease severity. Importantly, the data analysis, from raw video to ISI, is entirely automated, and does not require any human expert judgment or intervention. The clinician is directly given the ISI and corresponding interpretation band for a sample.

Our data show that the ISI interpretation bands correspond to the diagnosis established by retrospective physician adjudication, with sepsis or a non-infectious severe immune response diagnosed in nearly 90% of subjects in the Red Band but only 5% of subjects in the Green Band (Table 2). The ISI interpretation bands accurately identified subjects with more severe illness and in greater need of hospital resources as measured by SOFA score (maximum over 3 days), APACHE-II score, number of hospital-free days, and admission to the ICU (Fig 5). Moreover, the ISI interpretation bands clearly distinguished mortality risk in a high acuity population where all subjects presented to the ED with two or more criteria for SIRS and at least one sign of potential organ dysfunction (Fig 6A), and accurately identified patients with infections severe enough to cause morbidity and mortality (Fig 6B).

The ISI is designed for use in the ED and offers a number of distinct advantages over other approaches for sepsis diagnosis. The ISI can be measured from routinely collected blood samples and so there is no need for ED clinicians to order additional blood draws requiring specialized and expensive sample collection such as is needed for RNA profiling methods (18, 19). The run-time is short, targeted to less than 10 minutes end-to-end, including sample processing and image analysis (with the prototype unit used in this study, approximately 30 minutes). As such, ISI results can be available much sooner than other indicators of sepsis, enabling ED clinicians to intervene and potentially mitigate subsequent organ dysfunction. Unlike approaches reliant on electronic health records, the ISI does not require any patient or laboratory information. Another distinction is that the ISI can provide meaningful information at the point of treatment in the ED, whereas other approaches for sepsis diagnosis are intended for patient management in the ICU where clinical decisions identifying patients most at risk of clinical deterioration have already been made (20, 21). While some markers (e.g., lactate or C-reactive protein) provide an indication that organ damage has occurred, the ISI measures the activation of innate immunity that may lead to organ damage, thus providing an early warning. Lastly, because the test itself can be easily performed and interpreted, the ISI fits well within the ED workflow and may help improve the quality and efficiency of care.

This study has several limitations. First, the ISI was developed and internally validated with data from cohorts at two medical centers in Baton Rouge, which may limit the generalizability of the findings. For example, while we expect the age and sex distribution of the studies to be representative of what is expected of adults presenting to the ED across the US (when compared to data from the National ambulatory Medical Care Surveys collected by the Centers for Disease Control and Prevention), the cohorts in this study included a predominance of White and African American subjects and did not include significant numbers of subjects with Hispanic, Asian or other heritage. Additionally, the majority of subjects in the healthy study were White, and the race distribution of this cohort differed from that of the high and low acuity cohorts. Another limitation is that the study was not designed for longitudinal evaluation of the ISI or for direct head-to-head comparisons of the ISI with currently available biomarkers. Larger studies with different patient populations at multiple sites are currently underway to externally validate the performance of the ISI and to assess the potential clinical utility of the ISI in the ED context.

In conclusion, this study shows that the ISI has the potential to provide clinically actionable performance in distinguishing potentially septic patients. The ISI identifies patients with infections severe enough to cause morbidity and mortality, and can help ED clinicians in quickly recognizing sepsis and prioritizing hospital resources to care for those most at risk for clinical deterioration and death. Like other indicators of the host immune response, the ISI is pathogen agnostic and provides valuable information regardless of whether sepsis is caused by bacterial or viral infection. With a short turn-around time and no requirements for additional patient or laboratory data, the ISI is particularly well suited to aid ED clinicians in rapidly diagnosing sepsis.

## Acknowledgements

The authors would like to express their gratitude to the Office of Research at Our Lady of the Lake Regional Medical Center and the personnel for screening enrollment and overall study support: Jennifer Daigle, BSN; Katie M. Vance, Ph.D., Mandi Musso, Ph.D., Stephen Brierre, M.D., Tonya Jagneaux, M.D., Michael Sanchez, M.D., and Emilio Volz, M.D. The authors would also like to recognize the contributions of the team at Cytovale for their work in the set-up and completion of this study: Katherine Crawford, Sakshi Shah, Madeline Hem, Chris Dahlberg, Allison Walters, Mara Macdonald, Michael Samoszuk, Daniel Burgin, Peter Landwehr and Bliss Lambert.

## Supporting information

**S1 Fig. Flow chart for selection of evaluable subjects and exclusion of ineligible subjects.**

**S2 Fig. Receiver Operating Characteristic (ROC) curves showing the 3-fold cross-validation of the high acuity cohort.**

**S3 Fig. Distribution of cross-validated AUC for repeated 10-fold partitions of the high acuity cohort. Dotted black line indicates the mean AUC of 0.91.**

**S1 Table. Effect size, as measured by Cohen’s d, for data in Fig 2.**

**S2 Table. Baseline demographics (age, sex, and race) across interpretation bands for subjects in each of the three cohorts.**

**S1 Dataset. Spreadsheet including all data pertaining to the clinical characteristics, ISI underlying features, ISIs, and interpretation bands per subject for the three study cohorts.**

**S2 Dataset. Per cell data for aspect ratio, VEIR, and size for each of the 3 representative subjects presented in Fig. 1.**

