## Supporting Information for "Development and validation of a cellular host response test as an early diagnostic for sepsis"

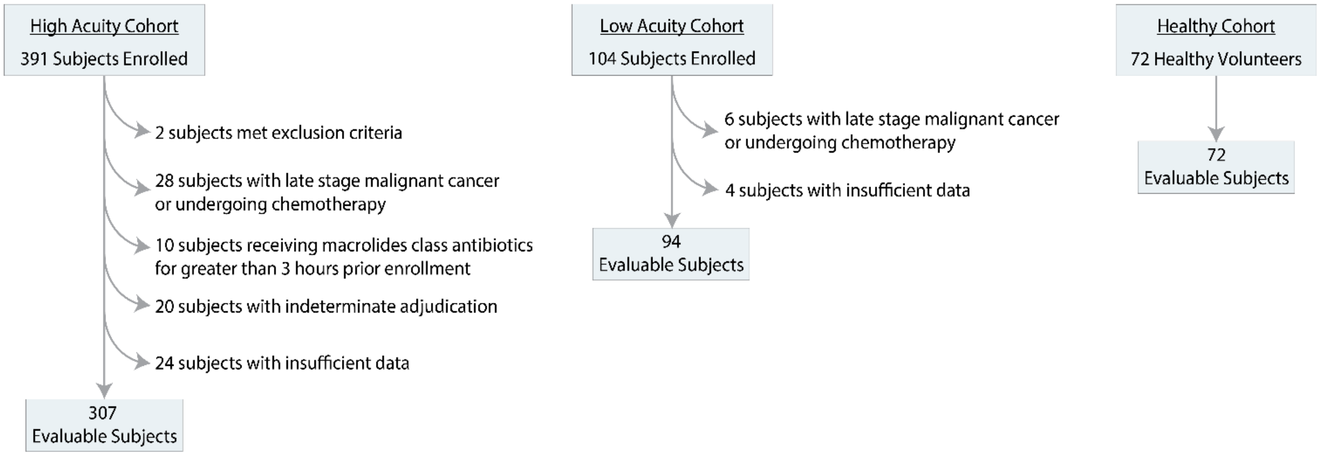


**S1 Fig. Flow chart for selection of evaluable subjects and exclusion of ineligible subjects.**

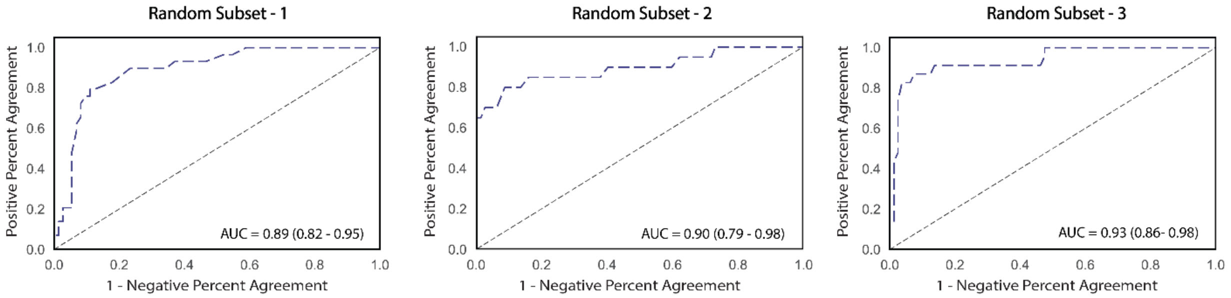


S2 Fig. Receiver Operating Characteristic (ROC) curves showing the 3-fold cross-validation of the high acuity cohort.

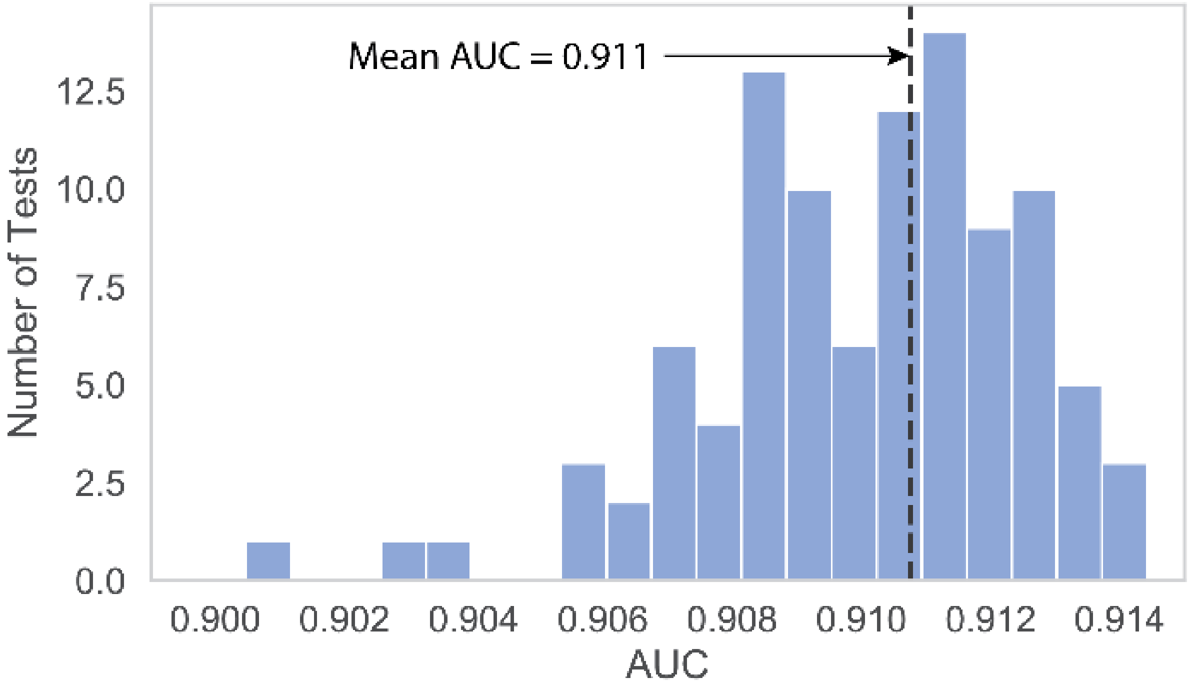


**S3 Fig. Distribution of cross-validated AUC for repeated 10-fold partitions of the high acuity cohort.** Dotted black line indicates the mean AUC of 0.91.

**S1 Table. Effect size, as measured by Cohen's d, for data in Fig 2.**

| **Cell type** | **Mean Aspect Ratio** | | | **Mean VEIR** | | |
|---|---|---|---|---|---|---|
| | **Healthy vs. SIRS 2+** | **SIRS 2+ vs. Septic** | **Healthy vs. Septic** | **Healthy vs. SIRS 2+** | **SIRS 2+ vs. Septic** | **Healthy vs. Septic** |
| Lymphocytes | 0.87 | -0.40 | 0.46 | -0.91 | 0.52 | -0.35 |
| Neutrophils | -1.16 | -0.68 | -1.99 | -0.71 | -1.43 | -1.90 |
| Monocytes | -0.22 | -0.54 | -0.77 | -0.79 | -1.42 | -2.50 |

VEIR, visco-elastic inertial response

**S2 Table. Baseline demographics (age, sex, and race) across interpretation bands for subjects in each of the three cohorts.**

| Characteristic | | Cohort | Green Band | Yellow Band | Red Band |
|---|---|---|---|---|---|
| Age, median (IQR) | | High Acuity | 58.7 (46-71) | 52.0 (63-77) | 57.0 (56-85) |
| Gender (Female), N (%) | | | 96 (49.2) | 29 (54.7) | 29 (49.2) |
| Race, N (%) | White | | 94 (48.2) | 40 (75.4) | 29 (49.2) |
| | African American | | 88 (45.1) | 10 (18.9) | 25 (42.4) |
| | Other | | 13 (6.7) | 3 (5.7) | 5 (8.5) |
| Age, median (IQR) | | Low Acuity | 54.4 (37-71) | 60.5 (51-64) | 61.9 (54-70) |
| Gender (Female), N (%) | | | 33 (45.2) | 4 (40.0) | 7 (63.6) |
| Race, N (%) | White | | 34 (46.6) | 6 (60.0) | 7 (63.6) |
| | African American | | 36 (49.3) | 4 (40.0) | 4 (36.4) |
| | Other | | 3 (4.1) | 0 (0.0) | 0 (0.0) |
| Age, median (IQR) | | Healthy | 52.5 (38-68) | 0 (0.0) | 0 (0.0) |
| Gender (Female), N (%) | | | 40 (55.6) | 0 (0.0) | 0 (0.0) |
| Race, N (%) | White | | 63 (87.5) | 0 (0.0) | 0 (0.0) |
| | African American | | 4 (5.6) | 0 (0.0) | 0 (0.0) |
| | Other | | 3 (4.2) | 0 (0.0) | 0 (0.0) |

Abbreviations: IQR, interquartile range (Q1 – Q3).
